# Evaluating Evo 2 for plant variant effect prediction

**DOI:** 10.64898/2026.03.24.714098

**Authors:** Yoshinobu Kato, Seiji Takayama, Sota Fujii

## Abstract

The genomic foundation model Evo 2 enables zero-shot variant effect prediction. Here, we evaluate its performance using *Arabidopsis thaliana* reproductive barrier genes with experimentally confirmed gain- and loss-of-function variants, and show that Evo 2 distinguishes functionally impactful variants. Together with a sign-reversal amplitude metric that recovers a variant missed by standard scoring, these results highlight the potential of Evo 2 for causal variant prioritization in plant GWAS and QTL mapping.

---

The emergence of large-scale genomic foundation models has opened new possibilities for predicting the functional consequences of genetic variants without task-specific training data. Evo 2 is a recently published biological foundation model trained on 9.3 trillion nucleotides from a curated genomic atlas spanning all domains of life, including plant genomes, with a context window of up to 1 million base pairs at single-nucleotide resolution^1^, representing one of the most comprehensive genomic sequence models available. Evo 2 enables zero-shot variant effect scoring: given a nucleotide sequence and a candidate variant, the model assigns a likelihood-based score reflecting the predicted impact on biological function^1^. This capability holds considerable promise for interpreting the functional significance of natural polymorphisms and for prioritizing candidate causal variants in genetic mapping studies.

Evo 2 has demonstrated strong predictive performance in initial benchmarks, including the prediction of functional impacts for clinically significant human variants^1^. This raises the prospect of Evo 2 serving as a powerful tool for prioritizing candidate causal variants within genomic regions identified by genome-wide association studies (GWAS) and quantitative trait locus (QTL) mapping — approaches that are central to both basic plant research and crop breeding programs^2,3^, yet are often limited by the difficulty of pinpointing the causal variant among many candidates. Recent plant DNA language models have begun to suggest that zero-shot scores may aid post-GWAS/QTL causal variant prioritization. PlantCaduceus, for example, combined zero-shot scoring with GWAS to identify the causal *su1* variant in sweet corn, but was developed as a plant-specific model pretrained on 16 angiosperm genomes^4^; here, we ask whether comparable utility extends to Evo 2, a general genomic foundation model, without plant-specific fine-tuning or additional training. Extending this characterization to encompass the full spectrum of functional variant classes (gain-of-function, loss-of-function, and near-neutral) within a single well-characterized biological system provides a particularly informative test of the model’s discriminative power. The most rigorous such tests require not only experimentally characterized variants of each functional class but also genes with contrasting selective histories and population-level phenotypic data reflecting the cumulative effects of multi-variant haplotypes. The plant interspecific reproductive barrier genes *STIGMATIC PRIVACY 1* (*SPRI1*) and *SPRI2* in *Arabidopsis thaliana* nicely satisfy these criteria. Both genes were originally identified through GWAS conducted across natural *A. thaliana* accessions^5,6^. Critically, the functional consequences of specific variants in both genes have since been established experimentally^5– 8^.

*SPRI1* encodes a pistil factor required for interspecific pollen rejection in self-incompatible relatives of *A. thaliana*^5^. Because *A. thaliana* is self-compatible^9^, the selective pressure maintaining SPRI1 function is greatly reduced, allowing a spectrum of functionally distinct variants, including loss-of-function alleles and a gain-of-function substitution, to accumulate in natural accessions^5^. To evaluate Evo 2 zero-shot variant effect prediction, we scored all natural variants at the *SPRI1* locus (Fig. 1a) catalogued in the 1001 Genomes Project^10^. For each variant, Δlikelihood was calculated as the change in log-likelihood from the wild-type (Col-0) to the variant sequence. Strongly negative Δlikelihoods, suggesting functional impact, were almost exclusively associated with variants in coding regions: gene-disrupting variants such as frameshifts, stop-gained, and splice-site changes accumulated in the negative tail, whereas variants in 5’UTR, 3’UTR, introns, and intergenic regions remained clustered near zero (Fig. 1b). On the other hand, variants with positive Δlikelihoods were considerably fewer, consistent with the expectation that strongly beneficial variants are rare, whereas most segregating variants are neutral or of small effect^11^. Among these variants, G155A (a glycine-to-alanine substitution at position 155), an experimentally confirmed gain-of-function variant carried by the *SPRI1A* haplotype^5,8^, showed a positive Δlikelihood, consistent with barrier enhancement. However, variants known to impair SPRI1 function, including E2V, N92H, and stop222C, carried by the *SPRI1B* haplotype^5,7^, also showed Δlikelihoods near zero or positive in the forward orientation (Fig. 1b), suggesting that forward-only scoring would fail to flag these variants as functionally significant.

**Fig. 1.**
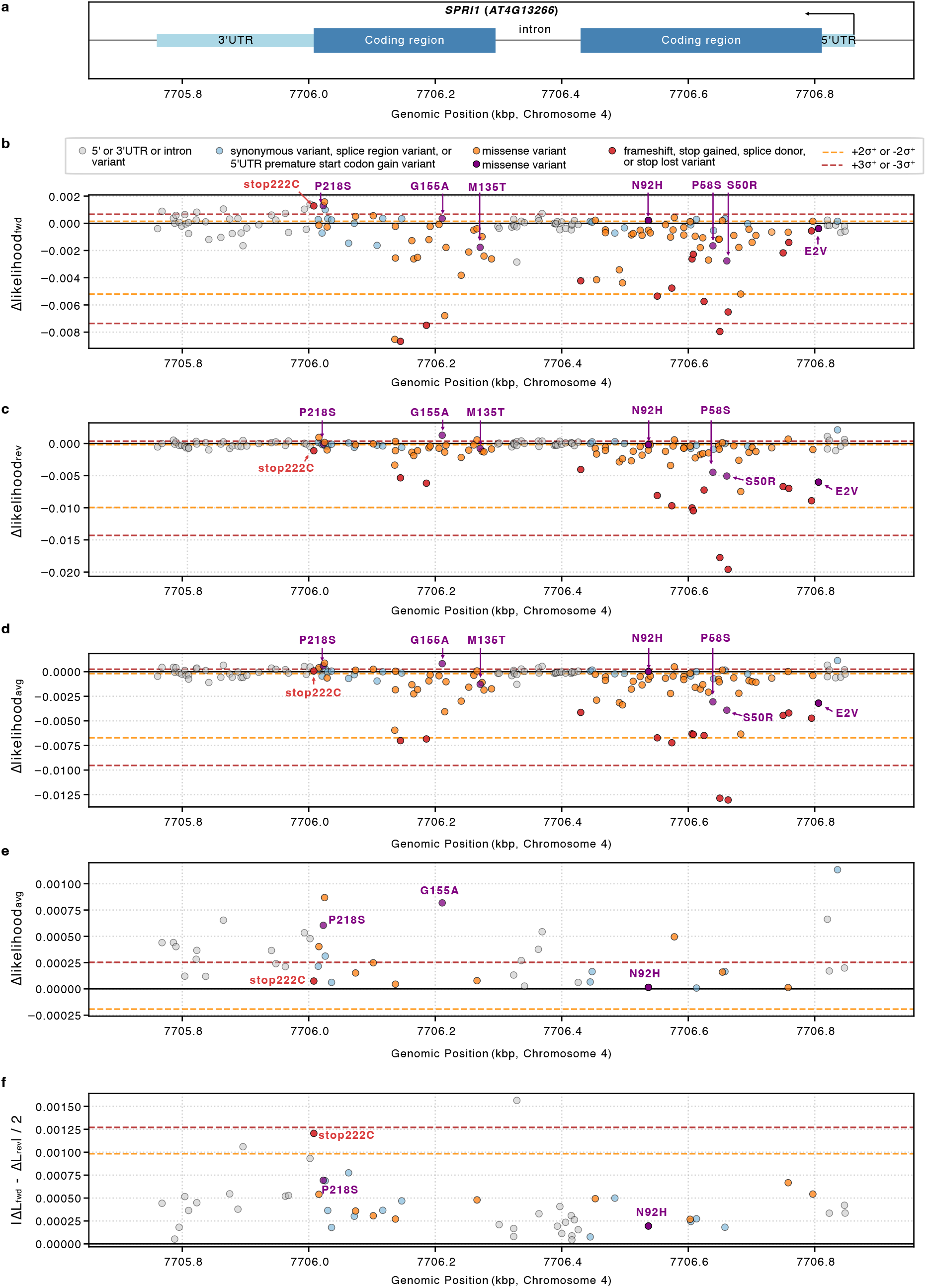
Evo 2 variant effect scores across the *SPRI1* locus. **(a**) Gene structure of *SPRI1*. The x-axis indicates the genomic position on chromosome 4 in kilobase pairs (kbp). **(b)** Forward-strand Δlikelihood (Δlikelihood_fwd_) of each variant relative to the Col-0 reference sequence. **(c)** Reverse-complement-strand Δlikelihood (Δlikelihood_rev_) of each variant relative to the Col-0 reference sequence. **(d)** The average score of Δlikelihood_fwd_ and Δlikelifood_rev_ (Δlikelihoodavg) of each variant relative to the Col-0 reference sequence. **(e)** Expanded view of panel (d), showing only variants with Δlikelihood_avg_ ≥ 0. **(f)** Sign-reversal amplitude, |Δlikelihood_fwd_ (ΔL_fwd_)− ΔL_rev_)| / 2, calculated only for variants in which Δlikelihood_fwd_ and Δlikelihood_rev_ carry opposite signs. Dashed lines indicate ±2σ (orange) and ±3σ (red) asymmetric guide lines. Source data is listed in Supplementary Table 1.

As reverse-complement inconsistency has been noted in DNA language models^12^, we also calculated Δlikelihoods in the reverse-complement orientation. (Fig. 1c). Among the gene-disrupting variants (frameshifts, stop-gained, etc.), scores were inconsistent between orientations: some variants that showed near-zero Δlikelihoods in the forward orientation yielded strongly negative values in the reverse complement orientation, while others that scored strongly negative in the forward orientation showed weaker values in the reverse (Fig. 1b, c). This orientation-dependent inconsistency suggests that single-orientation scoring may not reliably capture the functional severity of gene-disrupting variants. We therefore calculated an averaged Δlikelihood for each variant, as in the original Evo 2 study^1^, as the mean of the forward and reverse-complement scores (Fig. 1d). The averaged score placed frameshift and stop-gained variants consistently in the negative tail, except for stop222C. Although N92H remained near zero, E2V showed one of the most strongly negative averaged Δlikelihoods among missense variants (Fig. 1d).

Stop222C is a stop codon elimination caused by a frameshift at the 3′ boundary of the coding sequence, which extends the protein by 12 amino acids and reduces SPRI1 protein stability^7^. Although the averaged Δlikelihood was near zero due to cancellation of opposite-sign scores, the sign reversal itself may reflect an inherent functional ambiguity of stop codon loss and the resulting C-terminal extension: such additions can be stabilizing or destabilizing depending on the protein context, and the model’s directional disagreement may capture this uncertainty. In other proteins or contexts, analogous extensions can be beneficial. The model’s opposite verdicts in the forward and reverse orientations may thus reflect a genuine biological uncertainty about the functional consequence of this class of variant.

To flag such orientation-discordant variants systematically, we first selected variants showing sign reversal (i.e., forward and reverse Δlikelihoods with opposite signs) and ranked them by sign-reversal amplitude [|Δlikelihood_fwd_ − Δlikelihood_rev_| / 2]. The stop222C variant ranked among the top two by sign-reversal amplitude (Fig. 1f). Notably, this variant would have been missed by standard averaged Δlikelihood filtering alone, as its average approached zero; the sign-reversal amplitude metric thus serves as a complementary filter that specifically recovers this variant class that standard approaches fail to detect.

At the *SPRI2* locus (Extended Data Fig. 1a), the forward and reverse Δlikelihoods showed broadly similar distributions (Extended Data Fig. 1b, c), in contrast to the strand bias observed at *SPRI1* (Fig. 1b, c). The reduced strand bias in *SPRI2* may reflect its broader representation in the Evo 2 training corpus because the SPRI2 protein belongs to the SHI-family of zinc-finger transcription factors, a well-conserved protein family with numerous homologs across diverse plant lineages, likely providing abundant training signal in both orientations. The SPRI1 protein, by contrast, is restricted to a limited phylogenetic group within the Brassicaceae and lacks a well-characterized conserved domain. Therefore, relatively few homologous sequences were likely available during model training, potentially making the learned representations more sensitive to orientation. The overall scoring range in *SPRI2* was substantially narrower, and variants with strongly negative averaged Δlikelihoods were largely absent (Extended Data Fig. 1d). As SPRI2 directly controls cell wall xylan modification genes and indirectly upregulates *SPRI1* expression^6^, *SPRI2* is subject to broader functional constraints across tissues and developmental contexts, maintaining purifying selection in natural populations. Among variants with characterized functional outcomes, two substitutions in the C-terminal region (S343P and C345S) showed positive averaged Δlikelihoods (Extended Data Fig. 1d), consistent with their experimentally established gain-of-function effects on pollen rejection activity^6^.

To test whether Evo 2 captures functionally relevant variation at the population level, we calculated variant likelihood scores for natural *A. thaliana* haplotypes of *SPRI1* and compared them with phenotype scores from heterospecific pollen rejection assays. A negative correlation was observed between phenotype score and variant likelihood score (Fig. 2), such that haplotypes with greater pollen acceptance (lower SPRI1 function) tended to carry sequences predicted to be less optimal by Evo 2. Notably, no haplotypes with low variant log-likelihood scores (less than -7.520) were found in the phenotype score range of 1– 2.5, consistent with the requirement of intact SPRI1 function for the rejection of *M. littorea* pollen.

**Fig. 2.**
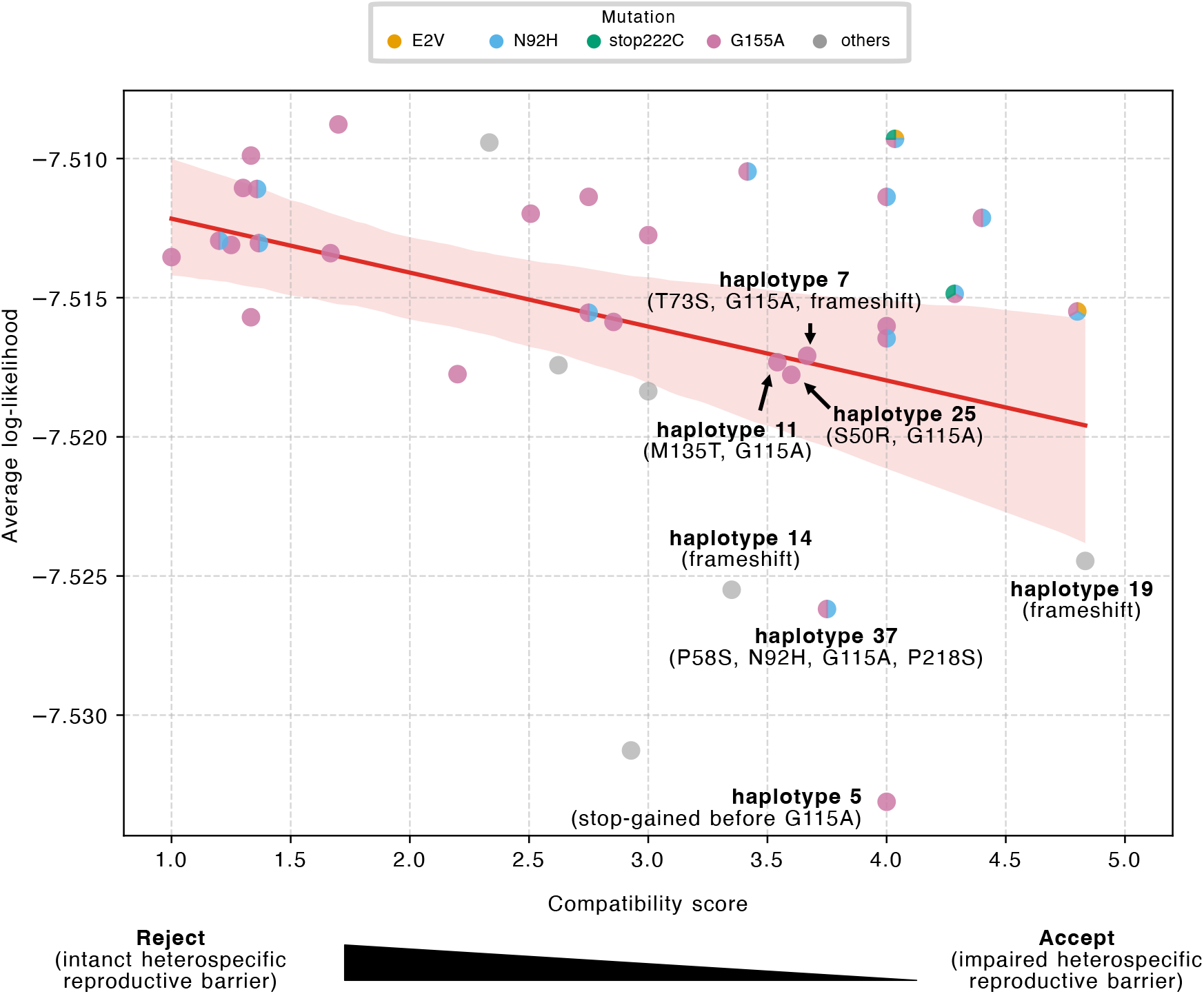
Correlation between Evo 2 variant likelihood scores and heterospecific (*M. littorea*) pollen rejection phenotype across natural *A. thaliana* haplotypes. Each point represents one *SPRI1* haplotype, plotted at the mean Evo 2 log-likelihood score and mean heterospecific pollen compatibility score for accessions sharing that haplotype, colored by the *SPRI1* variant it carries. The red line indicates a linear regression fit (95% confidence interval shaded; p = 0.024). Source data is listed in Supplementary Table 2. Compatibility score 1–5; higher scores reflect greater *Malcolmia littorea* pollen acceptance.

In contrast, some haplotypes, even though they carry the gain-of-function variant G155A, showed high compatibility scores (more than 3.5), suggesting reduced pollen rejection activity. Among haplotypes without frameshift or stop-gained variants, these correspond to haplotypes 11, 25, and 37, which carry G155A together with additional missense substitutions, and showed lower variant likelihood scores. The additional missense substitutions carried by these haplotypes — including S50R, P58S, M135T, H27N, and P218S — each show individual Δlikelihoods near zero or comparable to that of E2V, a variant that alone has little effect on SPRI1A function (Fig. 1d). Nevertheless, their co-occurrence with G155A partially overrides its gain-of-function effect, reducing pollen rejection activity — a functionally antagonistic combination that individual variant scoring alone would not readily predict.

Together, our results suggest that Evo 2 variant effect scoring offers a broadly applicable zero-shot framework for causal variant prioritization in plant genetics, both for prioritizing individual candidate variants within GWAS or QTL intervals and for capturing combinatorial effects through haplotype-level scoring, without the need for plant-specific fine-tuning or additional training.

## Materials and Methods

### Variant datasets

Natural variants at *SPRI1* and *SPRI2* loci were obtained from the 1001 Genomes Project ^10^. For each variant, the corresponding nucleotide substitution was individually introduced into the Col-0 reference genomic sequence to generate an 8,192 bp input sequence centered on the locus of interest. As both *SPRI1* and *SPRI2* are encoded on the reverse strand, input sequences represent the genomic (sense) strand, with the genes in reverse complement orientation relative to their coding sequences.

### Evo 2 variant effect scoring

Variant effect prediction was performed using the Evo 2 20B parameter model (version 0.5.0; Brixi et al., 2026). Variant effects were quantified as Δlikelihoods, calculated as the difference in log-likelihood between the variant and reference sequences: Δlikelihood = log P(variant) − log P(reference). Scoring was performed independently in both the forward (5’ to 3’) and reverse complement orientations, yielding two Δlikelihoods per variant. All scoring was zero-shot, with no task-specific fine-tuning of the model. For each variant, an averaged Δlikelihood was also calculated as Δlikelihood_avg_ = (Δlikelihood_fwd_ + Δlikelihood_rev_) / 2. For variants in which the forward and reverse Δlikelihoods carry opposite signs — indicative of orientation-discordant predictions — a sign-reversal amplitude was additionally calculated as |Δlikelihood_fwd_ − Δlikelihood_rev_| / 2. All scores were plotted against genomic position (kbp) to visualize the distribution of variant effects across each locus, with guide lines at ±2σ and ±3σ overlaid. To account for the characteristically skewed distributions of variant effect scores, these guide lines were calculated using asymmetric semi-standard deviations: σ^−^ was defined as the standard deviation of variants scoring below the distribution mean, and σ^+^ as that of variants scoring above the mean.

### Natural haplotype analysis

*SPRI1* sequences from natural *A. thaliana* accessions previously characterized ^5^ were used for haplotype-level scoring. For each accession, the full SPRI1 genomic sequence — spanning from the start codon to the stop codon and including an intron — was extracted and flanked by the corresponding Col-0 upstream and downstream sequences to generate an 8,192 bp input sequence. Each sequence was scored with the Evo 2 20B model in both the forward and reverse complement orientations, and the two log-likelihoods were averaged to yield a per-accession score. Heterospecific pollen compatibility scores were obtained from the previous study ^5^. Accessions were then classified into *SPRI1* haplotypes based on their variant composition, and both the Evo 2 score and the compatibility score were averaged within each haplotype. The resulting haplotype mean values were plotted.

## Acknowledgements

We thank M. Niidome., M. Saito, and S. Soneda for their technical assistance. This work was supported in part by Grants-in-Aid for Transformative Research Areas (22H05174 to S.F.), Grants-in-Aid for Scientific Research (21H05030 to S.T.; 24K01692 to S.F.), Grant-in-Aid for Challenging Exploratory Research (23K17987 to S.F.), Grant-in-Aid for Early-Career Scientists (23K14207 to Y.K.) from the Ministry of Education, Culture, Sports, Science and Technology of Japan (MEXT), and the Suntory Rising Stars Encouragement Program in Life Science (to S.F.).

## Author contributions

Y.K., S.T., and S.F. conceived the study and wrote the manuscript. Y.K. conducted Evo 2 analysis.

## Code Availability

The input data and custom Python scripts used in this study are available at Zenodo (DOI: 10.5281/zenodo.19201976).

## Legends of the Extemded Data Figure

**Extended Data Figure 1.**
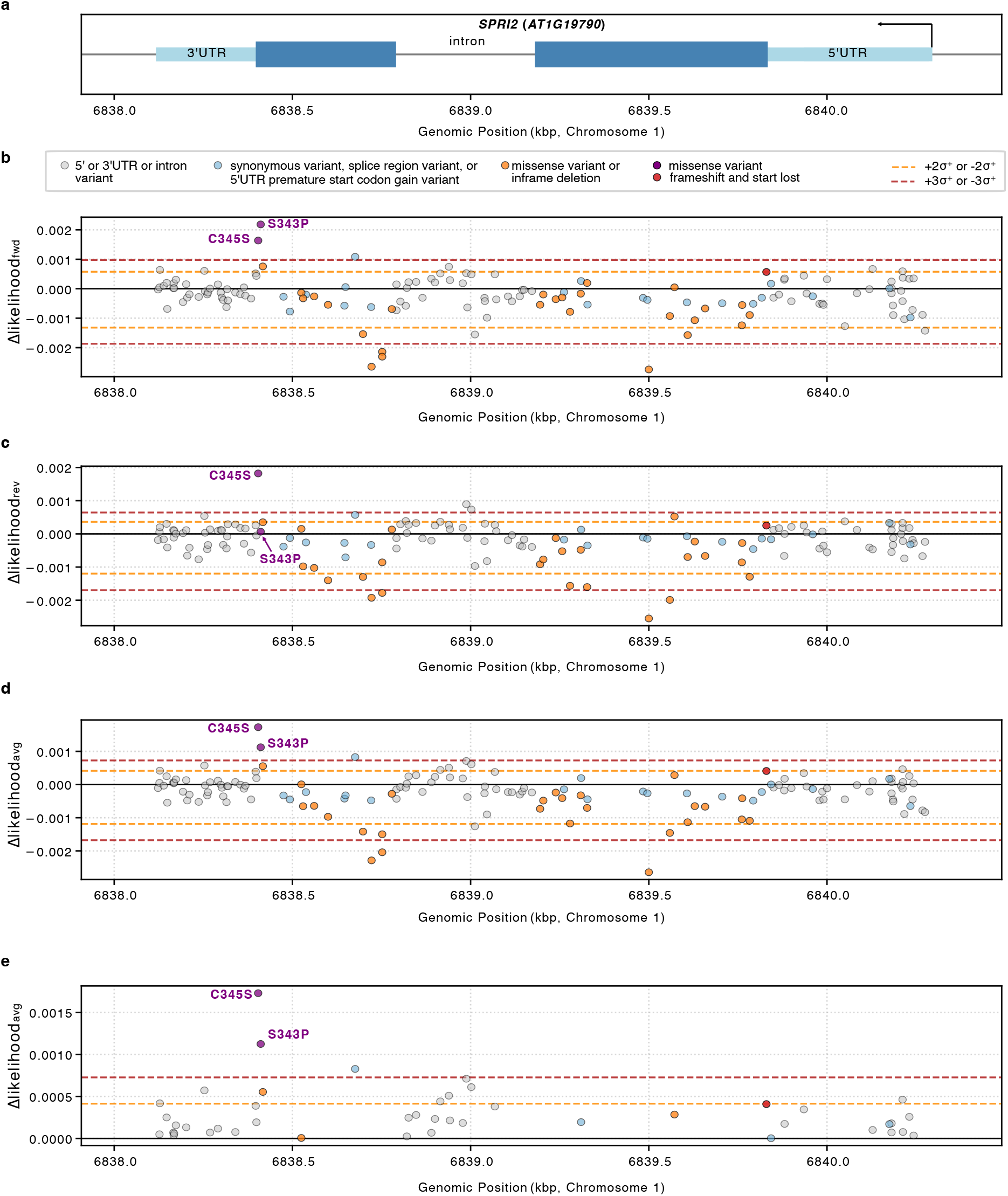
Evo 2 variant effect scores across the *SPRI2* locus. **(a**) Gene structure of *SPRI2*. The x-axis indicates the genomic position on chromosome 1 in kilobase pairs (kbp). **(b)** Forward-strand Δlikelihood (Δlikelihood_fwd_) of each variant relative to the Col-0 reference sequence. **(c)** Reverse-complement-strand Δlikelihood (Δlikelihood_rev_) of each variant relative to the Col-0 reference sequence. **(d)** The average score of Δlikelihood_fwd_ and Δlikelifood_rev_ of each variant relative to the Col-0 reference sequence. **(e)** Expanded view of panel (d), showing only variants with Δlikelihood_avg_ ≥ 0. Dashed lines indicate ±2σ (orange) and ±3σ (red) asymmetric guide lines. Source data is listed in Supplementary Table 3.

